# Dynamic Resting State Motor Network Connectivity of Neurotypical Children, the Groundwork for Network-Guided Therapy in Childhood Movement Disorders

**DOI:** 10.1101/2021.11.30.470606

**Authors:** Bethany L. Sussman, Sarah N. Wyckoff, Justin M. Fine, Jennifer Heim, Angus A. Wilfong, P. David Adelson, Michael C. Kruer, Varina L. Boerwinkle

## Abstract

**Background:** Normative childhood motor network resting-state fMRI effective connectivity is undefined, yet necessary for translatable dynamic resting-state network informed treatments in pediatric movement disorders.

**Method:** Cross-spectral dynamic causal modelling of resting-state fMRI was investigated in 19 neurotypically developing 5-7-year-old children. Fully connected six-node motor network models were created for each hemisphere including primary motor cortex, striatum, subthalamic nucleus, globus pallidus internus, thalamus, and contralateral cerebellum. Parametric Empirical Bayes with exhaustive Bayesian model reduction and Bayesian modeling averaging were used to create a group model for each hemisphere; Purdue Pegboard Test (PPBT) scores for relevant hand motor behavior were also entered as a covariate at the group level to determine the brain-behavior relationship.

**Results:** Overall, the resting-state functional MRI effective connectivity of motor cortico-basal ganglia-cerebellar networks was similar across hemispheres, with greater connectivity in the left hemisphere. The motor network effective connectivity relationships between the nodes were consistent and robust across subjects. Additionally, the PPBT score for each hand was positively correlated with the thalamus to contralateral cerebellum connection.

**Discussion:** The normative effective connectivity from resting-state functional MRI in children largely reflect the direction of inter-nodal signal predicted by other prior modalities and was consistent and robust across subjects, with differences from these prior task-dependent modalities that likely reflect the motor rest-action state during acquisition. Effective connectivity of the motor network was correlated with motor behavior, indicating effective connectivity brain-behavior relationship has physiological meaning in the normally developing. Thus, it may be helpful for future studies in children with movement disorders, wherein comparison to normative effective connectivity will be critical for network-targeted intervention.

**Impact Statement:** This is the first study to use pediatric resting-state functional MRI to create a normative effective connectivity model of the motor network and to also show correlation with behavior, which may have therapeutic implications for children with movement disorders.

## Introduction

One in 345 U.S. children have cerebral palsy (CP) (Winter et al., 2002), the most common movement disorder (MD) in childhood (Accardo & Capute, 2008), which has remain unchanged despite perinatal care intensification over the years (Reddihough & Collins, 2003). One in three with CP are severe and thus more likely to be medically resistant movement disorder (r-MD) with lifelong significant disability (Sadowska et al., 2020), which cost $1.3 million per patient (CDC 2003). Treatment of r-MD such as Parkinson’s disease (PD) in adults, via surgical interventions, is well established (Kleiner-Fisman et al., 2006; Liu et al., 2019). In children with dystonic CP, deep brain stimulation (DBS) placement began in 1996 (Coubes et al., 1999), and the most common targets in children are the globus pallidus internus (GPi) and the subthalamic nucleus (STN) (Larsh et al., 2021). According to a recent pediatric DBS meta-analysis, predictors of DBS response include shorter disease duration, idiopathic or inherited cause with negative MRI, truncal involvement, lower severity, and lack of skeletal deformity (Elkaim et al., 2019). The greater pediatric variety of MD causal pathogenesis yields higher variability in spatial and functional markers of dysfunctional networks, which may reduce surgical candidacy, increase surgical failure, and has prompted use of the invasive stereo-electroencephalogram (sEEG) approach to determine optimal targeting outcomes (Sanger et al., 2018). This technique resulted in unique target sites that otherwise would not have been considered and improved surgical outcomes from 50 to 80%.

In children with drug resistant epilepsy, non-invasive means of informing the sEEG placement by resting-state functional MRI (rs-fMRI) localizes the seizure onset zone with 90% correlation with sEEG, improves surgical outcomes, and increases surgical candidacy, thus may similarly inform DBS placement in MD (Boerwinkle et al., 2019b; oerwinkle, Foldes, et al., 2018; Boerwinkle et al., 2020; Boerwinkle et al., 2017; Chakraborty et al., 2020).

Advancing on these methods, effective connectivity (EC) of rs-fMRI measures appear to hold special promise in informing surgical targeting in childhood r-MD (cr-MD). Prior static connectivity (SC) measures of motor network (MN) configuration demonstrate parallel-interdependent loops, creating circuit redundancy and pathology resistance models (Alexander et al., 1986; Haber, 2003), and are associated with motor behavior across the lifespan into adulthood (Allen et al., 2011; Manza et al., 2015; Solé-Padullés et al., 2016). Unlike SC, EC incorporates time-dependent causal signaling between network locations (nodes). EC quantifies the nodal directed connectivity magnitude and polarity (i.e. excitatory or inhibitory), inferring a causal relationship. Thus, evaluation through EC-hypothesis-driven nodal-relationships and network configurations allows determination of which model, if any, explains the system signal variance best and locates the sources and sinks of atypical signal. Rs-fMRI EC in essential tremor (ET) detects different EC among MN nodes, even at rest (Park et al., 2017). Accordingly, EC predicts r-MD surgical outcomes in adults with ET (Park et al., 2017) and PD (Kahan et al., 2019; Kahan et al., 2014). However, this is not possible in individual cr-MD without an age-appropriate EC reference-standard, which is not reported to the authors’ knowledge.

While RSN are shown to be present as early as 28 weeks gestation (Doria et al., 2010), they also undergo changes associated with development. For example, the regions of the BG network shows SC increases associated with age, including connectivity with cortical regions (Solé-Padullés et al., 2016). The existence of age-related changes within SC networks supports the impetus to identify age-related changes in MN EC in typically developing children. Additionally, a recent study investigating lateralization of RSN SC in children reported age-related laterality increases (right putamen) and decreases (left putamen), leading to overall less lateralization with age. Other BG networks did not show lateralization changes with age (Agcaoglu et al., 2021). Therefore, we expect some hemispheric symmetry of EC BG networks in children, with potential asymmetry of connections involving the putamen. Thus, this study aims to determine normative MN EC in relation to motor behavior in children.

## Methods

The Institutional Review Board (IRB) of Phoenix Children’s Hospital (PCH) and National Institutes of Mental Health approved C-MIND data access (Cincinnati MR Imaging of NeuroDevelopment database https://cmind.research.cchmc.org/; see acknowledgements), which were acquired with informed consent. Both the initial data acquisition and current secondary analysis were completed in accordance with the Declaration of Helsinki’s ethical principles for medical research involving human subjects. The imaging and behavioral data used for this study are available through the National Institute of Mental Health Data Archive (NDA) through use a data use agreement between the NDA and qualifying institutions. Connectivity metadata for this study are available in Supplementary Files A and B.

### Sample

The rs-fMRI of 33 individual typically developing 5-7-year-old children were evaluated. Age, sex, T1-weighted (T1W), 5-minute rs-fMRI with ≤ 3 mm movement, and age-appropriate Purdue Pegboard right-hand dominance and motor skill score (PPBT) at the approximate time of scan were required. Those with history of self or family first generation neuropsychological history, no or left-hand preference, and lack of localizable region of interest anatomy were excluded. Of the 33 records evaluated, a total of 19 met study criteria.

### Acquisition and Preprocessing

A Philips 3T Achieva MRI, 32-channel head coil, and scan parameters: (1) T1W - standard inversion recovery prepared method, repetition time (TR) 8.1 msec, echo time (TE) 3.7 msec, flip angle (FA) 8°, shot interval 2800 msec, inversion time (TI) 939 msec, turbo field echo (TFE) shot duration/acquisition, 1851.9/1804.6 msec, field of view (FOV) 256 x 224 x 160 mm, voxel size 1 x 1 x 1 mm acquired in the sagittal plane with sensitivity encoding (SENSE) 5 min 15 secs; (2) rs-fMRI - TR 2000 msec, TE 35 msec, FA 90°, FOV 240 x 240 x 144 mm, matrix size 80 x 80, slice thickness of 4 mm, voxel size 3 x 3 x 4 mm, and 36 transverse slices of the whole brain. A standard rs-fMRI preprocessing pipeline (SPM12, www.fil.ion.ucl.ac.uk/spm, Wellcome Trust Centre for Neuroimaging, London, UK; (Friston et al., 1994) was utilized to eliminate the first 5 volumes, slice-timing correction, realignment, and linear coregistration of the echo planar imaging (EPI) data to the T1W, with no spatial smoothing. SPM12 and CAT12 (Gaser & Dahnke, 2016) were used to segment the T1W, which were resampled to 1 mm voxels and non-linearly registered to MNI space. CAT12 was then used to deform the relevant motor-related atlases (HMAT, (Mayka et al., 2006); BGHAT, (Prodoehl et al., 2008); and IBSR template, (Filipek et al., 1994)) to subject space for region of interest (ROI) specification.

#### General Linear Model

The GLM was first estimated over the whole brain to extract nuisance regressor time series from white matter (WM) and cerebrospinal fluid (CSF) spherical ROIs. A GLM was then re-estimated from the rs-fMRI data with a high-pass filter with a cut off frequency of 1/128s to remove low-frequency non-neural artifacts from data, the extra-cerebral time-series nuisance regressors (WM, CSF), and six nuisance regressors that captured head motion (Friston et al., 2003). Low-pass filtering (e.g. > 0.1 Hz) was not used because data in those frequencies may contain meaningful information in resting-state studies (Chen & Glover, 2015; Lin et al., 2015) and may be informative for comparison with clinical populations (Boerwinkle et al., 2017).

#### EC Analysis

The models were inverted using bilinear cross-spectral dynamic causal modeling (DCM) using DCM 12.5 (Friston et al., 2003; Friston et al., 2014). Six MN ROI per hemisphere were selected to incorporate key locations in the cortico-striato-pallido-thalamo-cortical and cerebello-thalamo-cortical pathways (Haber, 2003; Milardi et al., 2019; Quartarone et al., 2019) including: primary motor cortex (M1), striatum (STR), globus pallidus internus (GPi), subthalamic nucleus (STN), thalamus (THAL), and the contralateral cerebellum (CER). Each ROI location was identified from the predefined referenced atlases. Thus, a six-ROI (node) model was specified per hemisphere per subject (See Figure 1). The first eigenvariate of the rs-fMRI signal from the preprocessed and GLM-estimated six nodes were extracted. The first principal component eigenvariate was selected as it represented the dominant variation of activity in that node. Steady-state spectral amplitude and phase representations of each node’s activity were then obtained through Fourier transform. The local spectrum of each region was modeled as a power law distribution with an amplitude and scale, the latter indicating the frequency by amplitude slope (Friston et al., 2014). Thus, directional connectivity by DCM is achieved through EC estimated parameters of nodal frequency, auto- and cross-spectrum (herein also termed EC), between regions through multivariate auto-regression models of rs-fMRI blood oxygen derived (BOLD) signal.

**Figure 1.**
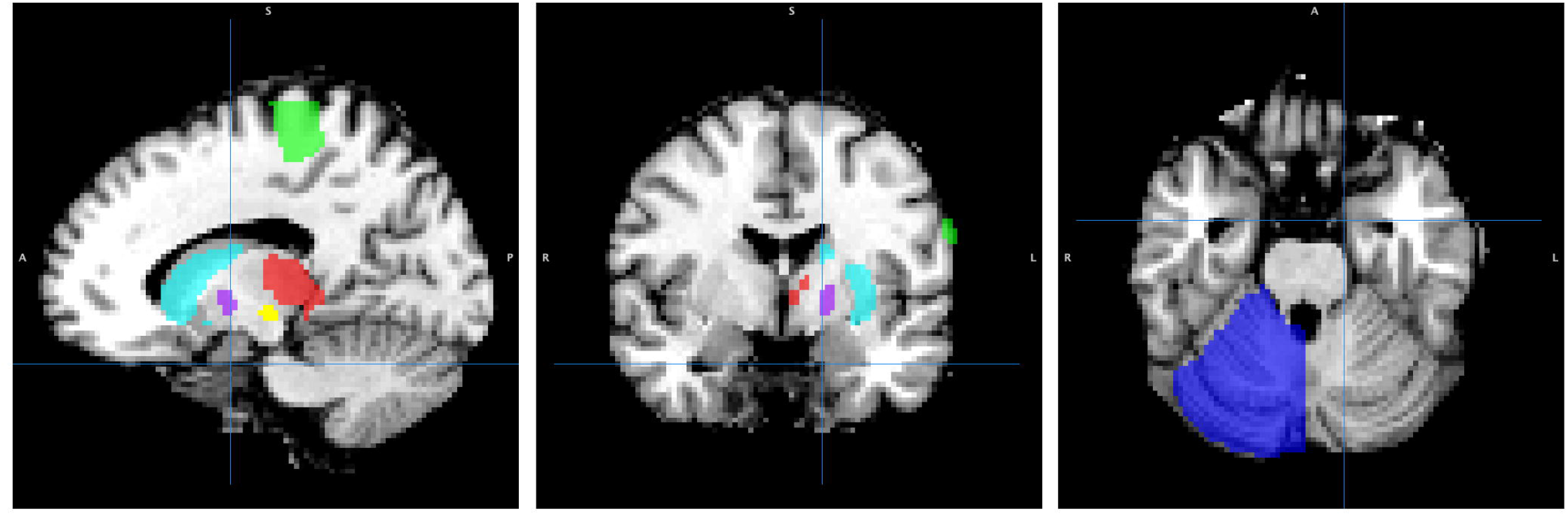
Regions of Interest. Sagittal, coronal, and axial views of the regions of interest in the left hemisphere model (radiological view). The right hemisphere model contains the same regions of interest with the opposite lateralization of each ROI. Key: Primary motor cortex (M1): green, striatum: cyan, subthalamic nucleus (STN): yellow, globus pallidus internus (GPi): purple, thalamus: red, contralateral cerebellum: blue. Atlases used: HMAT: M1; BGHAT: striatum (combination of caudate and putamen ROIs), STN, GPi; IBSR: thalamus, cerebellum). Figure is represented in MNI space, however, for analysis, ROIs were projected into subject anatomical space.

Variational Bayesian inversion was used to fit the differential equation connectivity model (Penny et al., 2011; Penny et al., 2010). This method of inversion involves fitting the DCM to maximize the likelihood of the model under prior EC parameter specifications. Model reduction or selection was not performed at the first level. The study protocol estimated models at the first level using an iterative empirical Bayes inversion scheme that alternated between estimating individual DCMs and estimating group effects to use as DCM priors. This method reduces the effects of local optima and draws subjects towards the group mean (Friston et al., 2015; Zeidman et al., 2019).

Second-level DCM with Parametric Empirical Bayes (PEB) was performed (Friston et al., 2015; Zeidman et al., 2019). PEB was used to create a group average for each hemisphere that preserved commonalities across the group **and differences associated with motor skill.** In addition to the group average, for each model the mean-centered raw Purdue Pegboard Test score (PPBT) nearest to the scan date from the contralateral hand was entered as a covariate. This was to investigate whether any EC parameters were associated with motor skill, herein termed ***<u>EC-PPBT to denote the brain-behavior association</u>***.

Because the goal was to create a proto-typical model of connectivity, it was assumed that all subjects would have the same model structure but may differ on EC parameter estimates. To maximize the similarities between subjects, parametric empirical Bayes framework was used for the group. PEB places greater weight on subjects and EC parameters that are less ‘noisy’ (Friston et al., 2015). Next, Bayesian Model Comparison (BMC) through Bayesian Model Reduction (BMR) (Friston et al., 2016) was performed on the full model space of each group hemisphere model, separately. BMR is a BMC method that effectively “prunes” redundant or uninformative EC parameters by exhaustively switching EC parameters and combinations of EC parameters of a fully connected and estimated model “on” and “off” and then efficiently computing the evidence and probability of each potential reduced model. BMR favors the simplest model that can explain the data (Friston et al., 2016). BMR was chosen over selecting specific models to compare to use the most data-driven model selection method in an essentially hypothesis-driven paradigm. Additionally, the investigators chose not to impose restrictions on connections that represent the direct, indirect, and hyperdirect pathways during model comparisons. The primary reason for this is that these pathways are based on knowledge of active paradigms (movement initiation, cessation, inhibition, etc.,) rather than an extended period of rest.

Through BMR, the similarities across the entire group were evaluated by exhaustive potential models and provided the optimized explanatory model. Thus, a Bayesian Model Average (BMA) was calculated over the 256 best models from the BMR to obtain the optimized model EC parameters by weighting their posterior probability (Friston et al., 2015; Penny et al., 2006). Further discussion is limited to EC parameters with posterior parameter estimates with evidence past the threshold of > 95% free energy modeled. EC parameter estimates with a probability > 0.95 underwent leave one out (LOO) cross-validation individually (**Figure 3**) to determine the effect sizes of the ***<u>EC-PPBT</u>*** correlations.

## Results

The median age for the 19 subjects (8 female) included was 6.3 years [IQR: 5.7-7.2]. The median righthand PPBT score was 9 [IQR: 9-11] and the median left-hand PPBT score was 8 [IQR: 7-9] (used for correlation to contralateral hemisphere). In one subject, the right STN activity could not be extracted due to small volume, and this subject was excluded from the right hemisphere model. Thus, 18 and 19 subjects’ data for the right and left hemisphere models respectively were utilized.

### Baseline Connectivity

The optimized mean EC parameters, passing BMR pruning and >95% probability, are delineated in **Figure 2A and 2B.** Detailed EC parameters and confidence intervals of all parameters can be found in **Supplementary Material A**.

**Figure 2.**
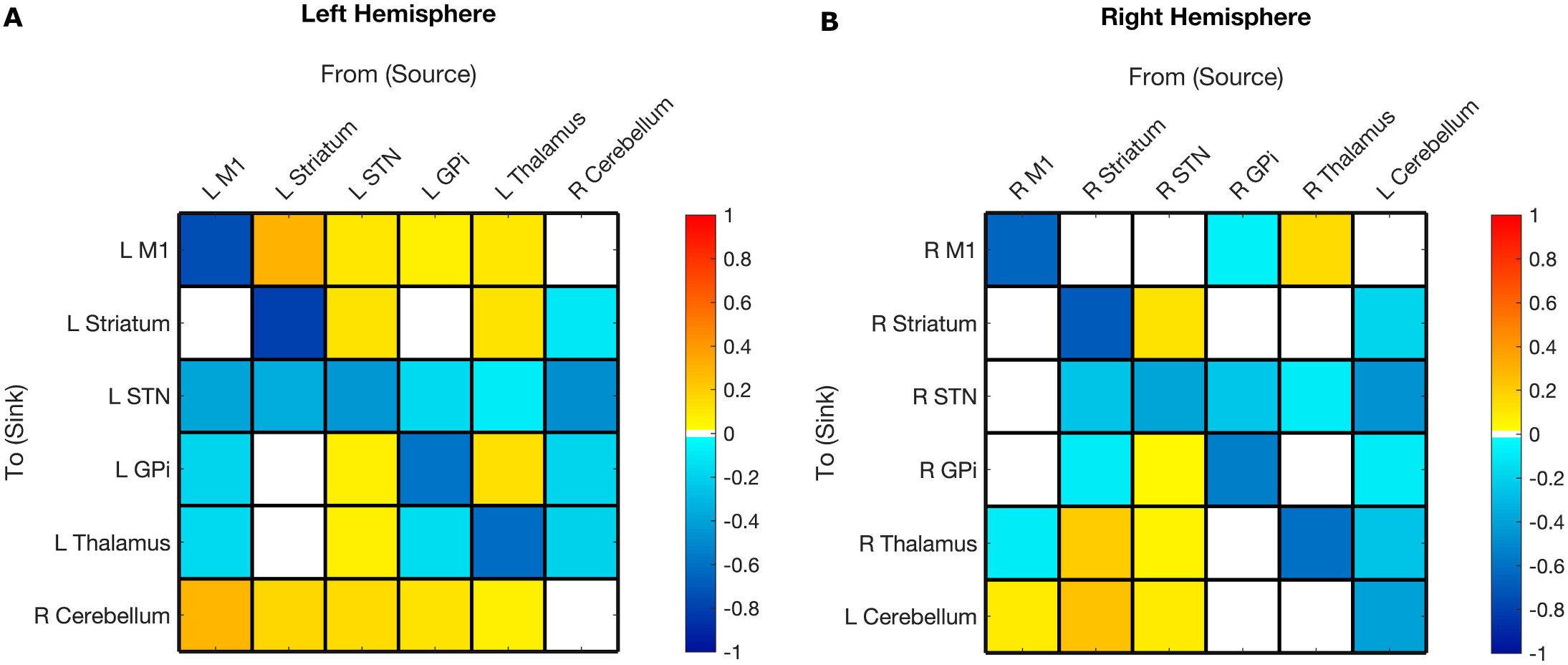
Hemispheric mean group effective connectivity. The y-axis lists the receiving nodes of the signal coming from the x-axis listed nodes. Blank squares are those that were pruned during Bayesian model reduction or did not meet the threshold for strong evidence of modulatory connection (> 0.95 posterior probability). The color bar spectrum indicates the normalized strength (magnitude) and the direction (color) of each connection in Hertz. Excitatory (positive) connections being ‘hot-red’ and inhibitory (negative) connections ‘cold-blue’. For example, the L striatum was excitatory toward the L M1, with greater magnitude than many other connections. Abbreviations: M1 – Primary Motor Cortex, STN – subthalamic nucleus, L – left, R – right. Note: Each hemisphere model features the contralateral cerebellum.

#### Hemispheric Symmetry

The polarity of the surviving connections was highly consistent across hemispheres. The only connection that was significant in both hemispheres and had opposite polarity was GPi➔M1. The left hemisphere had more overall connections (30 of 36 total potential connections) than the right hemisphere (24 of 36 total potential connections). Most of the surviving connections were the same across hemispheres. Pairwise correlation of the group BMA parameter estimates (Supplementary Materials 1A, 1B) showed a strong positive correlation between the two hemispheric models (r(34) = .77, *p* < 0.001; see Figure 3).

**Figure 3.**
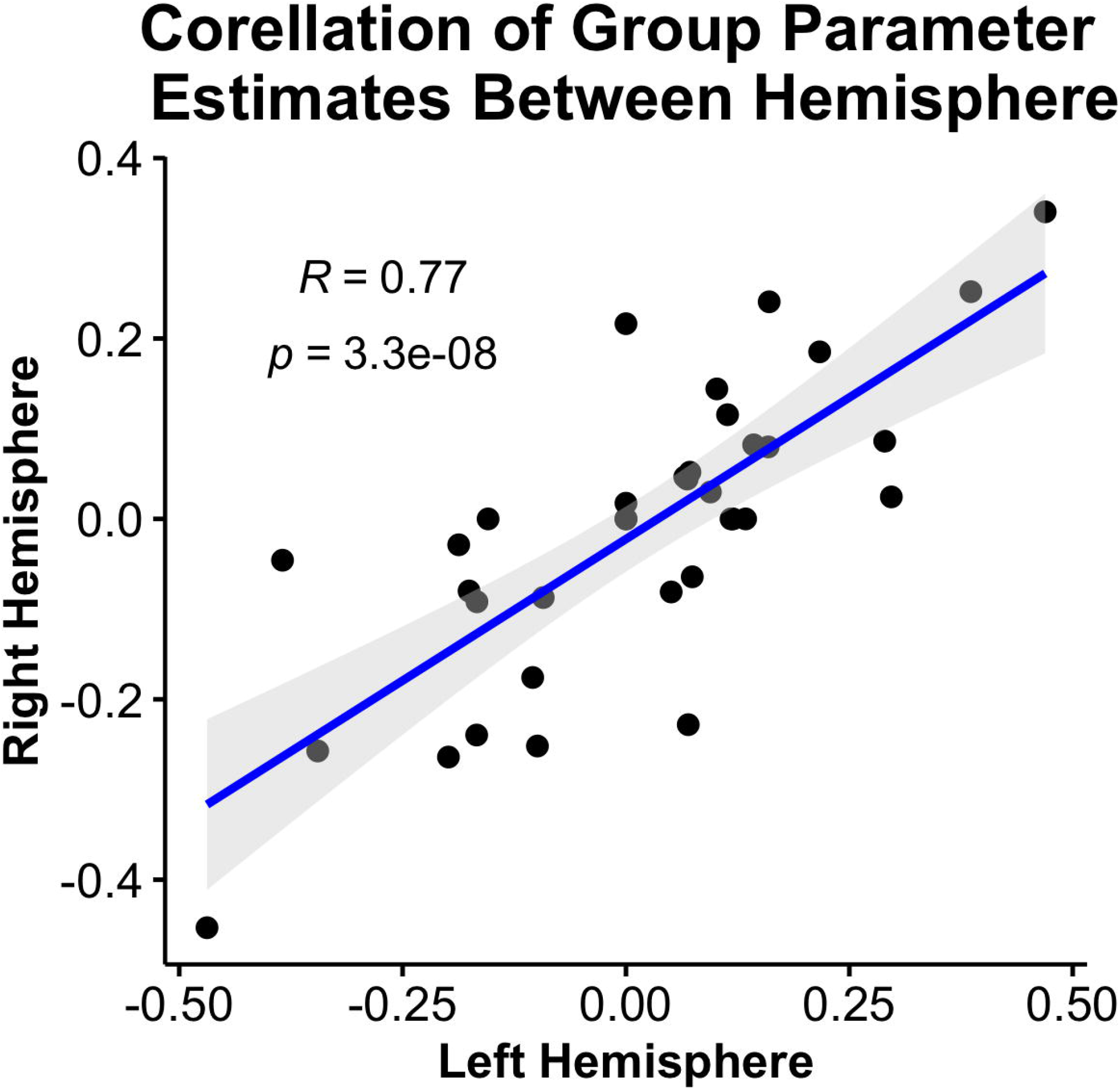
Correlation between hemisphere models of group BMA parameter estimates. The parameter estimate value for the left hemisphere is indicated on the x-axis, the parameter estimate value for the right hemisphere model is indicated along the y-axis. Each dot represents a connection in the model. The grey shaded region is the 95% confidence interval. All group BMA parameter estimates are included, regardless of posterior probability; pruned connections retain a zero value to be entered into the correlation. Pearson R (34) = 0.77, p < 0.001, indicating a strong positive correlation between the BMA group parameter estimates for the left and right hemisphere models.

#### Primary Motor

Modulations toward M1 were primarily excitatory with the only exception being rGPi➔rM1. This was also the only connection that had opposite polarity between hemispheres. Modulations from M1 were more mixed and excitatory toward the cerebellum but inhibitory toward deep grey locations. M1 had more surviving connections in the left hemisphere.

#### Striatum

Modulations toward striatum were primarily excitatory except those from the contralateral cerebellum. Modulations from striatum were mixed with positive modulations toward M1 in both hemispheres, cerebellum, and thalamus (right hemisphere), and negative modulations toward STN and GPi (right hemisphere).

#### Subthalamic Nucleus

Both hemisphere models showed inhibitory modulation toward the STN in most connections. In the left hemisphere, this included all connections. In the right hemisphere, this included all connections toward the STN except from M1.

#### Globus Pallidus Internus

Modulations toward GPi were mixed. Modulations from M1, striatum, and cerebellum were inhibitory while those from STN and thalamus were excitatory. Modulations from GPi were also mixed and more asymmetrical across hemispheres, with fewer connections from right hemisphere GPi.

#### Thalamus

All connections from thalamus survived in the left hemisphere, and all but one (STN) were excitatory. Fewer connections from thalamus survived in the right hemisphere, but the polarity of the surviving connections (M1, STN, thalamus) was the same as the left hemisphere.

#### Cerebellum (Contralateral)

In both the left and right hemisphere models, all connections from the contralateral cerebellum were inhibitory. All connections from the contralateral cerebellum to other nodes survived except right cerebellum to left M1 and left cerebellum to right GPi. All surviving modulations toward the contralateral cerebellum were excitatory. This included connections from all other nodes in the left hemisphere model and M1, striatum and STN in the right hemisphere model. All self-modulation survived except the right cerebellum self-inhibition (0.88 < 0.95).

### Motor Skill Correlation

The EC network parameters evaluated for the association with raw PPBT score with probability >0.95 are shown in Figure 4A and 4B. Detailed parameter estimates and confidence intervals of all connections with evidence can be found in **Supplementary Material B**.

**Figure 4.**
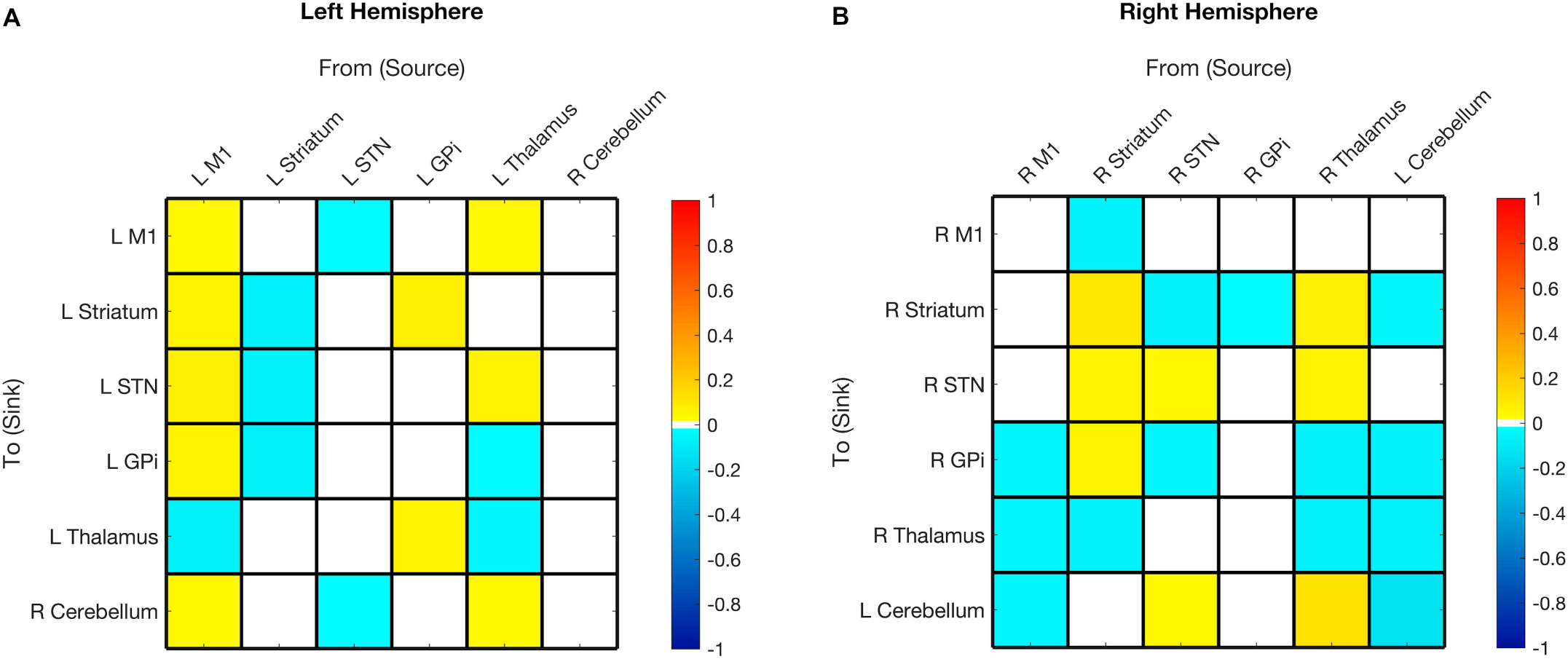
Hemispheric effective connectivity association with Purdue Pegboard Test score. The Purdue Pegboard Test score (PPBT) for the associated hand (i.e. right hand for left hemisphere, left hand for right hemisphere) was covaried with each queried connection. The color bar spectrum indicates the normalized strength and the direction of the relationship between a connection’s value and PPBT scores. For selfconnections, a positive parameter estimate on a self-connection (positive effect of a covariate) indicates a positive relationship between the covariate and the level of self-inhibition and a negative parameter estimate on a self-connection (negative effect of a covariate) indicates a negative relationship between the covariate and the level of self-inhibition. For example, the in left hemisphere self-connection for M1, higher PPBT scores were associated with more M1 self-inhibition, but, for the left hemisphere striatum self-connection, lower PPBT scores were associated with more self-inhibition. For all other connections, positive (hot) parameters indicate that more excitatory/less inhibitory connectivity values are associated with higher PPBT scores (positive relationship) and negative (cool) parameter values indicate that more inhibitory/less excitatory connectivity values are associated with higher PPBT scores (negative relationship). For example, the left hemisphere STN➔M1 connection, higher PPBT scores were associated with more inhibition, but for the left hemisphere Thalamus➔M1 connection, higher PPBT scores were associated with more excitation). Abbreviations: M1 – Primary Motor Cortex, STN – subthalamic nucleus, L – left, R – right. Note: each hemisphere model features the contralateral cerebellum.

### Leave-one-out cross validation

The EC-PPBT with strong evidence for brain-behavior relationship were relatively few and confidence intervals relatively wider compared to the estimates from the group average. The LOO cross-validation to determine the effect sizes of the correlations resulted in significant correlations for three remaining left hemisphere connections and five remaining right hemisphere connections. Significant connections are shown in Figure 5 and all LOO cross-validation results are listed in **Supplementary Table C** and are reported uncorrected for multiple comparisons due to the exploratory nature of this analysis.

**Figure 5.**
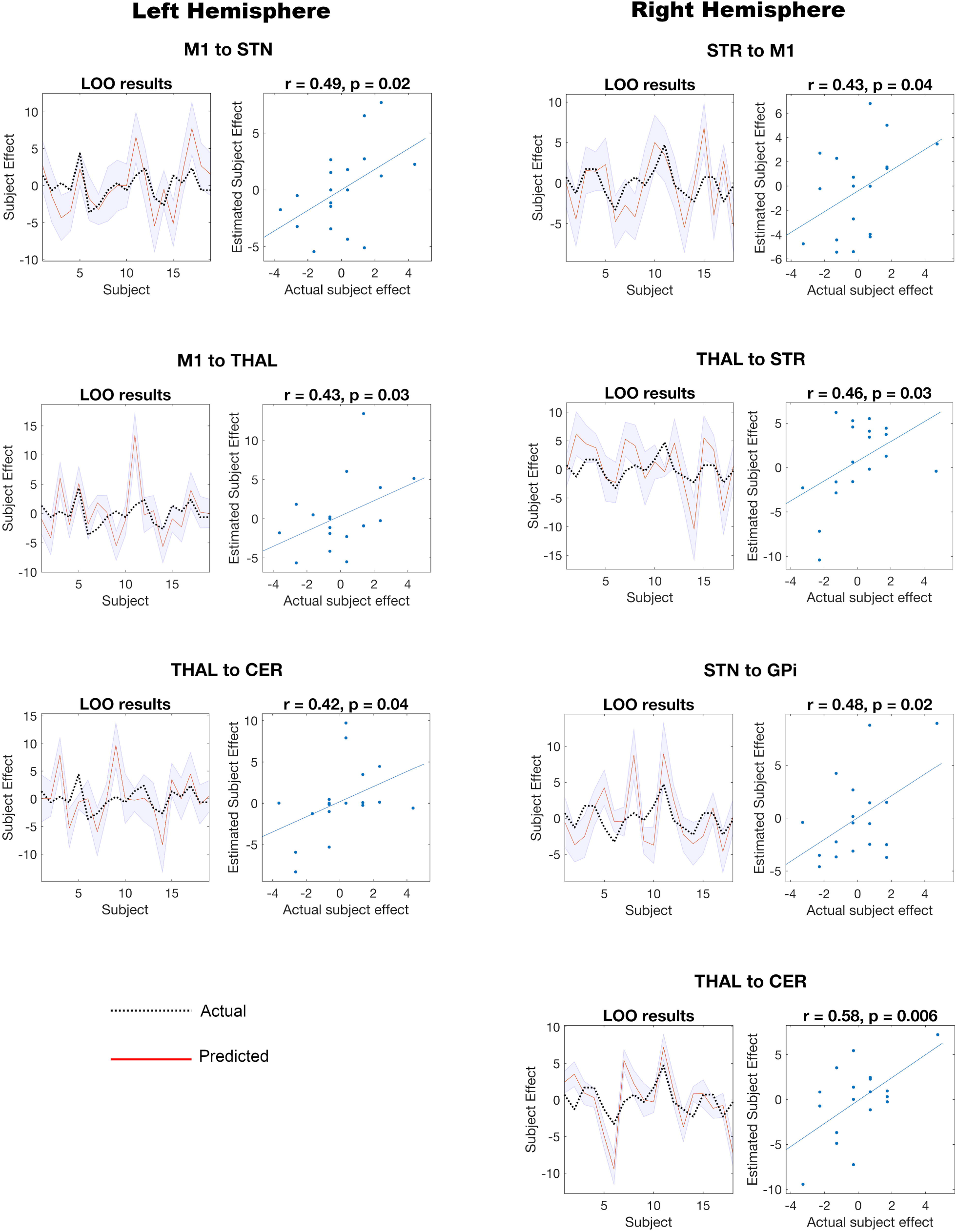
Consistency of Network Parameters to Motor Behavior. Results of the Purdue Pegboard Test score (PPBT) leave-one-out (LOO) cross-validation show the degree of consistency and significance of the network parameters (brain) to motor behavior, brain-behavior relationship. The individual LOO cross-validation tests are organized by hemisphere and connection. LOO cross-validation is performed by iteratively removing each subject’s data and, predicting their data given the rest of the subjects’ data, and then comparing their estimated value to the actual value for that subject. For each LOO cross-validation test, there are two plots. **Left:** The line graph for each comparison (right), shows the out-of-samples estimate of the mean-centered PPBT (red line) for each subject plus variance (purple shaded area, 90% CI) and the actual group effect (black dotted line). Thus, the purple envelope is relatively close to the predicted estimate and the predicted estimate is relatively close to the actual estimate. The narrower the purple envelope, the higher the degree of confidence and consistency the brain behavior relationship of the given connection. **Right:** The blue dotted scatter plot compares the actual subject effect to the estimated subject effect of the PPBT brain-behavior relationship for each subject using an out-of-samples Pearson correlation coefficient (r). Although the brain-behavior correlations were in the 0.42-0.58 range, those listed were significant with p<0.05 (uncorrected). Left and right hemisphere degrees of freedom were 17 and 16, respectively. Only significant tests are listed here. All tested correlations and values are listed in **Supplementary Table C**.

## Discussion

This is the first report of normative pediatric EC of the MN, which is necessary for meaningful individual comparisons in future cr-MD studies. The expected model validating benchmarks are: (1) consistency and robustness of results and correlation with motor behavior in normative data, (2) consistency across hemispheres, with expected differences for given handedness, and (3) future study determination of expected differences in those with focal network pathology also correlating with behavior.

### Motor Skill

Although several EC parameters were correlated with PPBT in the PEB analysis, the confidence intervals were relatively large, with only a very few connections surviving LOO cross-validation in effect size determination. None-the-less this significant brain-behavior relationship with relatively small variability in normal motor behavior, in a narrow age range, and relatively small sample size, is indicative of this measure’s potential in comparing subjects with gross pathology to normal. This is consistent with (Kahan et al., 2014) who showed that clinical score changes after STN-DBS for PD were also associated with changes in EC.

The EC-PPBT THAL➔CER was the only common brain-behavior relationship between hemispheres, which was more excitatory with higher PPBT scores. This is consistent with several prior modality studies indicating the cerebello-thalamo-cortical pathway’s role in skilled movement in primates (Horne & Butler, 1995). Additionally, fractional anisotropy of the denato-thalamic-cortical tract from left dentate nucleus to right DLPFC is associated with earlier rhythm-related finger motor skill (Schulz et al., 2014). Clinically, DBS (Koller et al., 2000) and thalamotomy (Chen et al., 2006) of the region of the thalamus that receives cerebellar input improves the adaptive control of reaching in essential tremor. This suggests that the integrity of cerebellar output toward the thalamus is important for the adaptive control of reaching behavior (Chen et al., 2006), which is used in tasks such as PPBT. While the significant correlations were THAL➔CER, and not the opposite (the thalamus does not have a known output to the cerebellum that is not mediated by cortex), this may be due to the resting-state condition or representative of an indirect connection.

It is also important to consider that PPBT scores were, in general, not correlated with the same parameters for each hemisphere. This implies that motor dexterity in dominant versus non-dominant hand are differentially associated with EC in the relevant hemispheric motor pathways. This is something that should be taken into consideration when describing the relationship between clinical scores and potential neuroimaging biomarkers.

### Subthalamic Nucleus

The importance of the STN in suppressing unwanted movements (Milardi et al., 2019) in children likely awake but instructed to not move, is reflected herein where the STN EC connections were the most consistent across subjects and numerous to other MN nodes. All connections to and from STN were inhibitory and excitatory, respectively. This degree of consistency and robustness also suggests a high degree of STN MN role conservation, the reproducibility of the EC by DCM, and reflects clinical experience targeting the STN in deep brain stimulation (DBS) for PD and dystonia (Lumsden et al., 2017).

In prior models, excitation *to* the STN in movement suppression is expected. However, inhibition was found, possibly because the net circuitry inhibition resulted in an overall effect of all signal into the STN being negative. Supportively, an optogenic fMRI study in mice, using spectral DCM to compare BG circuit function when stimulating D1 versus D2 dopamine pathways, consistently saw either negative or no modulation from cortex to STN (Bernal-Casas et al., 2017).

All connections *from* the STN in right and left hemispheres were excitatory, consistent with prior research (Haber, 2003). Given that many of these connections are not direct, this speaks toward the capacity of EC to measure dynamic effects across intermediate nodes. Some connections, for example STN➔THAL, may be expected to be inhibitory during a task due to the nature of the indirect pathway. However, the net excitation may be due to resting conditions and the fact that we applied no model constraint for specific connections to replicate pathways known from task-activity.

### Globus Pallidus Internus

The GPi disperses the final output of the BG neuromodulatory signal to the thalamus. The GPi’s primary modulators are the STN and the striatum (Albin et al., 1989; Haber, 2003). The GPi is likely key to dystonia and it is the most frequent DBS target for its treatment (Lumsden et al., 2017). Yet DBS to the GPi effectiveness varies widely, indicating that in some, initiation of dystonic signal may be from cerebellar or cortical networks (Neychev et al., 2011; Poston & Eidelberg, 2012; Quartarone et al., 2019; Rothkirch et al., 2018) or that other deep grey structures may contribute to dystonic signal.

Aligned with prior indirect pathway connections, herein an increase in GPi activity led to decrement in thalamic activity. Also, an increase of STN activity led to increase of GPi activity in the left hemisphere (Albin et al., 1989). Although the right hemisphere GPi➔THAL connection was pruned, the GPi had an inhibitory effect on M1, likely reflecting this indirect connection. Thus, with such a robust expected ‘typical’ profile of MN connections, individual connectivity biomarkers specific to pathophysiology location, may be predictive of outcome.

In a local field potential-MEG study of patients with dystonia, pallido-cerebellar coherence in the alpha band (7-13Hz) correlated negatively with dystonic symptom severity in adult patients with segmental or cervical dystonia (Neumann et al., 2015). Interestingly, herein connectivity between the GPi and the contralateral cerebellum remained in both directions for the left hemisphere model, but only CER➔GPi (inhibitory) in the right hemisphere model. Even with iterative BMR, which favors the simplest model possible, the models with the highest evidence still included nodes in the cerebellum and M1. This suggests that *both* are important when considering typical BG function. This finding is aligned with more recent models of cerebello-basal ganglia-thalamo-cortical pathways (Caligiore et al., 2017; Quartarone et al., 2019). Also, virus tracing studies have indicated an STN output that targets the dentate nucleus (Bostan et al., 2010) and a cerebellar target to STR (Hoshi et al., 2005).

### Primary Motor

In prior models, M1 signal is largely to the STN and striatum and the major input to M1 is the thalamus (Albin et al., 1989; Alexander et al., 1986; Haber, 2003). Many models do not include M1 and CER (Battistella & Simonyan, 2019; Jahfari et al., 2011; Kahan et al., 2014; Rowe et al., 2010), thus less is understood about this connection (see Dirkx et al., 2016; Kahan et al., 2019; Rothkirch et al., 2018 for models with cerebellum). The present models do not show CER➔M1, only an excitatory modulation from each M1 to the contralateral CER. It is unclear why the contralateral CER➔M1 modulations are not present in these results. Even before BMR and BMA, CER➔M1 was not present for either hemisphere in the group PEB analysis.

In a recent task-fMRI DCM study comparing healthy controls and patients with writer’s cramp dystonia during a finger tapping task, M1 to cerebellum connectivity was more excitatory in control than patients with writer’s cramp and connectivity from cerebellum to M1 was more excitatory for the patients than controls. The cerebellum was also more excitatory toward the putamen for patients than controls (Rothkirch et al., 2018). A recent study in PD showed winning resting-state DCM models with bidirectional modulations between the cerebellum and M1 that were not modulated by DBS. However, model comparison did not include switching modulations between M1 and cerebellum on and off (Kahan et al., 2019). The difference in polarity for outgoing connections from the cerebellum in the existing literature and current study may be due to task vs. resting-state or differences in model specification and reduction.

### Thalamus

The thalamus primary output is excitatory to the cerebral cortex, however, it also projects directly to the striatum (Haber & Calzavara, 2009). Here, in the left hemisphere model (dominant), the thalamus was highly influential toward other structures, but only excitatory toward M1 in the right hemisphere. Furthermore, when comparing this to associations with PPBT scores, in the left hemisphere model higher PPBT scores were associated with increased excitatory signal for THAL➔STR. This is consistent with the finding that thalamostriatal projections are associated with skilled initiation of movement sequences in mice (Díaz-Hernández et al., 2018). It is possible that, with enough increased skill, the right-hemisphere THAL➔STR parameter would not be pruned, mirroring the left hemisphere model. The thalamus had several incoming connections in the final model for both hemispheres. Interestingly, each hemisphere had four incoming connections, but there appears to be a trade-off between an inhibitory connection from GPi (left) and an excitatory connection from striatum (right). Taken together, these findings suggest a trade off in connectivity patterns associated with both hand dominance and motor skill.

### Cerebellum (Contralateral)

In both hemispheres, modulations from the contralateral cerebellum were inhibitory. Neuronal tracing studies in both rodents and non-human primates support connections between the cerebellum and BG and have demonstrated evidence of cerebellar projections toward BG primarily targeting the ‘indirect’ pathway (Hoshi et al., 2005). Findings from a virus-tracing neuroanatomical study in non-human primates indicates that cerebellar output targets the indirect pathway of the BG, specifically via a disynaptic pathway through the intralaminar thalamic nuclei to the putamen (Hoshi et al., 2005).

In both hemisphere models, the cerebellum had an inhibitory effect on thalamus and striatum, demonstrating a directed (effective) functional connection. There is also electrophysiological evidence from mice that that simultaneous stimulation of cortical and cerebellar inputs to the striatum result in long-term potentiation of cortico-striatal loops, whereas simulation of cortical inputs alone induces long-term depression to the same (Chen et al., 2014). Additionally, there has been an increasing amount of evidence for cerebellar contribution to movement disorders such as dystonia. Rodent models have shown dystonic movement as a result of cerebellar lesions (Neychev et al., 2011) and neurophysiological studies in humans show evidence that the cerebellum is involved in the pathophysiology of dystonia, though no conclusive evidence as the site of origin (Shakkottai et al., 2017).

### Hemispheric Symmetry

The left hemisphere had more connections, in line with expectations from hemispheric dominance. A recent study of handedness and EC in motor systems showed that right-handed subjects had stronger effective coupling between more key left-hemisphere regions than right during unimanual movement, and left-handed subjects showed less hemispheric asymmetry overall (Pool et al., 2014). As the subjects in the current study were right-hand dominant, it is possible that this contributed to more coupling in the left hemisphere. Further evaluation comparing EC in left hand dominant individuals may be needed for similar comparisons in this populations.

In general, connections were similar for each hemisphere with more left hemisphere connections. Overall, the value of connection in one hemisphere was strongly correlated with the value of that connection in the other hemisphere (Figure 3). This was true whether the pruned (BMA) group estimates or the PEB group estimates were used for the corellation. However, the striatum was one of the only regions for which outgoing modulations were more populated in the right hemisphere. Striatal modulations toward STN and GPi were inhibitory and striatal modulations toward non-deep grey regions (as well as thalamus) were excitatory.

The only connection that conflicted in polarity between hemispheres was from GPi➔M1, which was excitatory in the left hemisphere and inhibitory in the right hemisphere. Like the hemispheric discrepancies in the striatum, it is possible that there is a tradeoff between the number and location of connections, or the number of connections and polarity of connections at a given location. Since the GPi projects out from the BG to the thalamus and ultimately to the motor cortex, and its outputs have been associated with movement disorders (Milardi et al., 2019; Neychev et al., 2011), the polarity difference for GPi➔M1 between hemispheres is potentially of importance in motor control. However, this parameter was not correlated with motor skill.

### Model Comparisons

This EC MN model has several fundamental differences compared to prior works, thus differences are expected and highlighted. (1) The EC model includes ***<u>more subcortical ROIs</u>*** than some MN models (Battistella et al., 2019; Diekhoff-Krebs et al., 2017; Kahan et al., 2014; Zhang et al., 2020), because a more global view of the MN dynamics was desired, and rs-fMRI did not impose relevant spatial limits. (2) Unlike most MN studies conducted under active or (Jahfari et al., 2011; Rothkirch et al., 2018) passive task (e.g. stimulation, tremor onset; (Bernal-Casas et al., 2017; Dirkx et al., 2016; Kahan et al., 2014) conditions, the subjects herein were instructed to ***<u>rest without motion</u>*** for 5 minutes, theoretically requiring either continuous impulse inhibition and/or a state of executively sanctioned relaxation. (3) In 5-7-year-old children told to be perfectly still, it is possible that this may result in “***<u>active” stillness</u>**,* wherein instead of relaxing, intermittent stiffening or any number of difficult to visualize actions to maintain stillness are possible. This could have consequences on the network dynamics. (4) The direct, indirect (Albin et al., 1989), and hyperdirect (Nambu et al., 2002) pathways were not explicitly modeled (e.g. Bernal-Casas et al., 2017; Dirkx et al., 2016; Jahfari et al., 2011; Kahan et al., 2014). Instead, this EC model makes ***<u>no assumptions on which nodes are effectively connected, nor how</u>***. This model is based on the relatively well-established premise that when two nodes are not immediately connected, they could still have an influence on each other. This ***<u>more data-driven</u>*** facet of the EC model was intentional because the conditions of the data acquisition had no prior established precedent. Thus, novel network dynamics of this condition may be discovered. (5) Prior models largely did not include query for ***<u>bidirectional signal</u>*** (e.g. both forward and backward directed connections; (Dirkx et al., 2016; Kahan et al., 2014), whereas the study EC model did make this allowance. This is key, because if signal from A to B is the only direction queried, then B to A will remain unknown. Including feedback or feedforward connections may change the results of the system dynamics. It may be argued that only previously established connections should be modeled, however within the non-independent parallel loop system of the MN all such EC are biologically plausible. Further as a loop, all members activity should or could have direct or downstream result on all other members. (6) Developmentally related EC MN differences compared to adult are unknown, thus any portion of the differences herein could be attributed to age. (7) Many prior models only investigated a single hemisphere or did not explicitly consider hemispheric differences (Dirkx et al., 2016; Kahan et al., 2014; Rothkirch et al., 2018), thus differences in direction of network signal are less well established.

#### Limitations

Since this study only includes right-handed children, generalizations to left-handed children should be made cautiously. The data in this dataset also contain relatively short rs-fMRI sessions, which carries a higher concern for noise. At the time the data were analyzed, the C-MIND database was the only publicly available dataset with healthy, neurotypical, young children that also included hand dominance and motor scores. Future studies may expand to larger age spread to build developmental trajectory, include lefthanded participants to identify any differences, and compare clinical groups, such as children with movement disorders who can be completed under conscious sedation with surgically reliable signal acquisition in children (Boerwinkle et al., 2019a; Boerwinkle, Foldes, et al., 2018; Boerwinkle, Vedantam, et al., 2018).

## Conclusions

In this study, we used spectral DCM with a PEB approach to identify a pattern of typical directed connectivity in healthy 5-7-year-olds in a motor network consisting of cortical, deep grey, and cerebellar locations as well as associations with motor skill. BMR and BMA led to similar models of intrinsic connectivity for both hemispheres, even when each hemisphere was separately specified, estimated, and reduced – providing evidence of reliability for this technique. Leave-one out cross-validation showed that the left and right hemisphere model parameters had primarily unique associations with motor-skill as measured by PPBT score but shared a relationship with the THAL➔CER connection. While prior task-based fMRI methods allow for a more direct testing of established motor action pathways, rs-fMRI is often more plausible to collect in children, particularly those with complex movement disorders. Establishing models of normative directed connectivity in a resting-state has the potential to improve precision for therapeutic treatments in complex movement disorders.

## Acknowledgements

Data used in the preparation of this article were obtained from the C-MIND Data Repository created by the C-MIND study of Normal Brain Development. This is a multisite, longitudinal study of typically developing children from ages newborn through young adulthood conducted by Cincinnati Children’s Hospital Medical Center and UCLA and supported by the National Institute of Child Health and Human Development (Contract #s HHSN275200900018C). A listing of the participating sites and a complete listing of the study investigators can be found at https://research.cchmc.org/c-mind. Data were released from the NIMH Data Archive (NDA) in 2020.

This manuscript reflects the views of the authors and may not reflect the opinions or views of the NIH.

## Authorship Confirmation Statement

Conception/design of work – All authors; Analysis – BLS, VLB, JMF; Interpretation – BLS, VLB, MCK; Drafting – BLS, SNW, VLB, MCK; Critical Revision for important intellectual content – All authors; Approval of final version – All authors; Agreement to be accountable for all aspects of the work in ensuring that questions related to the accuracy or integrity of any part of the work are appropriately investigated and resolved. – All authors

## Disclosure Statement

None of the authors have any disclosures.

## Funding Statement

This work was supported in part by a grant from the University of Arizona/Valley Research Partnership (PI: Boerwinkle) Grant: P2-4014.

## Notes

### Competing Interest Statement

The authors have declared no competing interest.

## References

Accardo PJ, & Capute AJ. (2008). Capute & Accardo’s Neurodevelopmental Disabilities in Infancy and Childhood: Neurodevelopmental diagnosis and treatment (Vol. 1): Brookes Pub.

Agcaoglu O, Muetzel RL, Rashid B, et al. Lateralization of Resting-State Networks in Children: Association with Age, Sex, Handedness, Intelligence Quotient, and Behavior. Brain Connect 2021. doi:10.1089/brain.2020.0863

Albin RL, Young AB, & Penney JB. The functional anatomy of basal ganglia disorders. Trends in Neurosciences 1989, 12:10. doi:10.1016/0166-2236(89)90074-X

Alexander GE, DeLong MR, & Strick PL. Parallel organization of functionally segregated circuits linking basal ganglia and cortex. Annual review of neuroscience 1986, 9:1. 357-381.

Allen EA, Erhardt EB, Damaraju E, et al. A baseline for the multivariate comparison of resting-state networks. Frontiers in systems neuroscience 2011, 5. doi:10.3389/fnsys.2011.00002

Battistella G, & Simonyan K. Top-down alteration of functional connectivity within the sensorimotor network in focal dystonia. Neurology 2019, 92:16. doi:10.1212/wnl.0000000000007317

Bernal-Casas D, Lee HJ, Weitz AJ, et al. Studying Brain Circuit Function with Dynamic Causal Modeling for Optogenetic fMRI. Neuron 2017, 93:3. doi:10.1016/j.neuron.2016.12.035

Boerwinkle VL, Cediel EG, Mirea L, et al. Network Targeted Approach and Postoperative Resting State Functional MRI are Associated with Seizure Outcome. Ann Neurol 2019a. doi:10.1002/ana.25547

Boerwinkle VL, Cediel EG, Mirea L, et al. Network Targeted Approach and Postoperative Resting State Functional MRI are Associated with Seizure Outcome. Ann Neurol 2019b. doi:10.1002/ana.25547

Boerwinkle VL, Foldes ST, Torrisi SJ, et al. Subcentimeter epilepsy surgery targets by resting state functional magnetic resonance imaging can improve outcomes in hypothalamic hamartoma. Epilepsia 2018, 59:12. doi:10.1111/epi.14583

Boerwinkle VL, Mirea L, Gaillard WD, et al. Resting-state functional MRI connectivity impact on epilepsy surgery plan and surgical candidacy: prospective clinical work. J Neurosurg Pediatr 2020. doi:10.3171/2020.1.Peds19695

Boerwinkle VL, Mohanty D, Foldes ST, et al. Correlating Resting-State Functional Magnetic Resonance Imaging Connectivity by Independent Component Analysis-Based Epileptogenic Zones with Intracranial Electroencephalogram Localized Seizure Onset Zones and Surgical Outcomes in Prospective Pediatric Intractable Epilepsy Study. Brain Connect 2017, 7:7. doi:10.1089/brain.2016.0479

Boerwinkle VL, Vedantam A, Lam S, et al. Connectivity changes after laser ablation: Resting-state fMRI. Epilepsy Res 2018, 142. doi:10.1016/j.eplepsyres.2017.09.015

Bostan AC, Dum RP, & Strick PL. The basal ganglia communicate with the cerebellum. Proceedings of the National Academy of Sciences 2010, 107:18. doi:10.1073/pnas.1000496107

Caligiore D, Pezzulo G, Baldassarre G, et al. Consensus Paper: Towards a Systems-Level View of Cerebellar Function: the Interplay Between Cerebellum, Basal Ganglia, and Cortex. Cerebellum 2017, 16:1. doi:10.1007/s12311-016-0763-3

Chakraborty AR, Almeida NC, Prather KY, et al. Resting-state functional magnetic resonance imaging with independent component analysis for presurgical seizure onset zone localization: A systematic review and meta-analysis. Epilepsia 2020. doi:10.1111/epi.16637

Chen CH, Fremont R, Arteaga-Bracho EE, et al. Short latency cerebellar modulation of the basal ganglia. Nature Neuroscience 2014, 17:12. doi:10.1038/nn.3868

Chen H, Hua SE, Smith MA, et al. Effects of human cerebellar thalamus disruption on adaptive control of reaching. Cerebral cortex (New York, N.Y.: 1991) 2006, 16:10. doi:10.1093/cercor/bhj087

Chen JE, & Glover GH. BOLD fractional contribution to resting-state functional connectivity above 0.1 Hz. Neuroimage 2015, 107. doi:10.1016/j.neuroimage.2014.12.012

Coubes P, Echenne B, Roubertie A, et al. [Treatment of early-onset generalized dystonia by chronic bilateral stimulation of the internal globus pallidus. Apropos of a case]. Neurochirurgie 1999, 45:2. 139-144.

Díaz-Hernández E, Contreras-López R, Sánchez-Fuentes A, et al. The Thalamostriatal Projections Contribute to the Initiation and Execution of a Sequence of Movements. Neuron 2018, 100:3. doi:https://doi.org/10.1016/j.neuron.2018.09.052

Diekhoff-Krebs S, Pool E-M, Sarfeld A-S, et al. Interindividual differences in motor network connectivity and behavioral response to iTBS in stroke patients. NeuroImage. Clinical 2017, 15. doi:10.1016/j.nicl.2017.06.006

Dirkx MF, den Ouden H, Aarts E, et al. The Cerebral Network of Parkinson’s Tremor: An Effective Connectivity fMRI Study. The Journal of Neuroscience 2016, 36:19. doi:10.1523/JNEUROSCI.3634-15.2016

Doria V, Beckmann CF, Arichi T, et al. Emergence of resting state networks in the preterm human brain. Proc Natl Acad Sci U S A 2010, 107:46. doi:10.1073/pnas.1007921107

Elkaim LM, Alotaibi NM, Sigal A, et al. Deep brain stimulation for pediatric dystonia: a meta-analysis with individual participant data. Dev Med Child Neurol 2019, 61:1. doi:10.1111/dmcn.14063

Filipek PA, Richelme C, Kennedy DN, et al. The young adult human brain: an MRI-based morphometric analysis. Cereb Cortex 1994, 4:4. doi:10.1093/cercor/4.4.344

Friston KJ, Harrison L, & Penny WD. Dynamic causal modelling. NeuroImage 2003, 19:4. doi:10.1016/S1053-8119(03)00202-7

Friston KJ, Holmes AP, Worsley KJ, et al. Statistical parametric maps in functional imaging: a general linear approach. Human brain mapping 1994, 2:4. 189-210.

Friston KJ, Kahan J, Biswal B, et al. A DCM for resting state fMRI. Neuroimage 2014, 94:100. doi:10.1016/j.neuroimage.2013.12.009

Friston KJ, Litvak V, Oswal A, et al. Bayesian model reduction and empirical Bayes for group (DCM) studies. NeuroImage 2016, 128. doi:10.1016/j.neuroimage.2015.11.015

Friston KJ, Zeidman P, & Litvak V. Empirical Bayes for DCM: A Group Inversion Scheme. Frontiers in Systems Neuroscience 2015, 9:164. doi:10.3389/fnsys.2015.00164

Gaser C, & Dahnke R. CAT-a computational anatomy toolbox for the analysis of structural MRI data. Hbm 2016, 2016. 336–348.

Haber SN. The primate basal ganglia: parallel and integrative networks. Journal of Chemical Neuroanatomy 2003, 26:4. doi:10.1016/j.jchemneu.2003.10.003

Haber SN, & Calzavara R. The cortico-basal ganglia integrative network: the role of the thalamus. Brain Res Bull 2009, 78:2–3. doi:10.1016/j.brainresbull.2008.09.013

Horne MK, & Butler EG. The role of the cerebello-thalamo-cortical pathway in skilled movement. Prog Neurobiol 1995, 46:2–3. 199-213.

Hoshi E, Tremblay L, Féger J, et al. The cerebellum communicates with the basal ganglia. Nature neuroscience 2005, 8:11. 1491-1493.

Jahfari S, Waldorp L, van den Wildenberg WPM, et al. Effective Connectivity Reveals Important Roles for Both the Hyperdirect (Fronto-Subthalamic) and the Indirect (Fronto-Striatal-Pallidal) Fronto-Basal Ganglia Pathways during Response Inhibition. The Journal of Neuroscience 2011, 31:18. doi:10.1523/JNEUROSCI.5253-10.2011

Kahan J, Mancini L, Flandin G, et al. Deep brain stimulation has state-dependent effects on motor connectivity in Parkinson’s disease. Brain 2019, 142:8. doi:10.1093/brain/awz164

Kahan J, Urner M, Moran R, et al. Resting state functional MRI in Parkinson’s disease: the impact of deep brain stimulation on ‘effective’ connectivity. Brain 2014, 137:4. doi:10.1093/brain/awu027

Kleiner-Fisman G, Herzog J, Fisman DN, et al. Subthalamic nucleus deep brain stimulation: summary and meta-analysis of outcomes. Mov Disord 2006, 21 Suppl 14. doi:10.1002/mds.20962

Koller WC, Pahwa PR, Lyons KE, et al. Deep brain stimulation of the Vim nucleus of the thalamus for the treatment of tremor. Neurology 2000, 55:12 Suppl 6. S29–33.

Larsh T, Wu SW, Vadivelu S, et al. Deep Brain Stimulation for Pediatric Dystonia. Seminars in Pediatric Neurology 2021, 38. doi:10.1016/j.spen.2021.100896

Lin F-H, Chu Y-H, Hsu Y-C, et al. Significant feed-forward connectivity revealed by high frequency components of BOLD fMRI signals. NeuroImage (Orlando, Fla.) 2015, 121. doi:10.1016/j.neuroimage.2015.07.036

Liu Y, Li F, Luo H, et al. Improvement of Deep Brain Stimulation in Dyskinesia in Parkinson’s Disease: A Meta-Analysis. Frontiers in Neurology 2019, 10:151. doi:10.3389/fneur.2019.00151

Lumsden DE, Kaminska M, Ashkan K, et al. Deep brain stimulation for childhood dystonia: Is ‘where’ as important as in ‘whom’? Eur J Paediatr Neurol 2017, 21:1. doi:10.1016/j.ejpn.2016.10.002

Manza P, Zhang S, Hu S, et al. The effects of age on resting state functional connectivity of the basal ganglia from young to middle adulthood. Neuroimage 2015, 107. doi:10.1016/j.neuroimage.2014.12.016

Mayka MA, Corcos DM, Leurgans SE, et al. Three-dimensional locations and boundaries of motor and premotor cortices as defined by functional brain imaging: A meta-analysis. NeuroImage 2006, 31:4. doi:10.1016/j.neuroimage.2006.02.004

Milardi D, Quartarone A, Bramanti A, et al. The Cortico-Basal Ganglia-Cerebellar Network: Past, Present and Future Perspectives. Frontiers in Systems Neuroscience 2019, 13:61. doi:10.3389/fnsys.2019.00061

Nambu A, Tokuno H, & Takada M. Functional significance of the cortico–subthalamo–pallidal ‘hyperdirect’ pathway. Neuroscience research 2002, 43:2. doi:10.1016/S0168-0102(02)00027-5

Neumann W-J, Jha A, Bock A, et al. Cortico-pallidal oscillatory connectivity in patients with dystonia. Brain 2015, 138:7. doi:10.1093/brain/awv109

Neychev VK, Gross RE, Lehéricy S, et al. The functional neuroanatomy of dystonia. Neurobiology of disease 2011, 42:2. doi:10.1016/j.nbd.2011.01.026

Park HJ, Pae C, Friston KJ, et al. Hierarchical Dynamic Causal Modeling of Resting-State fMRI Reveals Longitudinal Changes in Effective Connectivity in the Motor System after Thalamotomy for Essential Tremor. Front Neurol 2017, 8. doi:10.3389/fneur.2017.00346

Penny WD, Friston KJ, Ashburner JT, et al. (2011). Statistical parametric mapping: the analysis of functional brain images: Elsevier.

Penny WD, Mattout J, & Trujillo-Barreto N. Bayesian model selection and averaging. Statistical Parametric Mapping: The analysis of functional brain images. London: Elsevier 2006.

Penny WD, Stephan KE, Daunizeau J, et al. Comparing families of dynamic causal models. PLoS Comput Biol 2010, 6:3. doi:10.1371/journal.pcbi.1000709

Pool E-M, Rehme AK, Fink GR, et al. Handedness and effective connectivity of the motor system. NeuroImage 2014, 99. doi:10.1016/j.neuroimage.2014.05.048

Poston KL, & Eidelberg D. Functional brain networks and abnormal connectivity in the movement disorders. Neuroimage 2012, 62:4. doi:10.1016/j.neuroimage.2011.12.021

Prodoehl J, Yu H, Little DM, et al. Region of interest template for the human basal ganglia: comparing EPI and standardized space approaches. Neuroimage 2008, 39:3. doi:10.1016/j.neuroimage.2007.09.027

Quartarone A, Cacciola A, Milardi D, et al. New insights into cortico-basal-cerebellar connectome: clinical and physiological considerations. Brain 2019, 143:2. doi:10.1093/brain/awz310

Reddihough DS, & Collins KJ. The epidemiology and causes of cerebral palsy. Aust J Physiother 2003, 49:1. doi:10.1016/s0004-9514(14)60183-5

Rothkirch I, Granert O, Knutzen A, et al. Dynamic causal modeling revealed dysfunctional effective connectivity in both, the cortico-basal-ganglia and the cerebello-cortical motor network in writers’ cramp. NeuroImage: Clinical 2018, 18. doi:10.1016/j.nicl.2018.01.015

Rowe JB, Hughes LE, Barker RA, et al. Dynamic causal modelling of effective connectivity from fMRI: Are results reproducible and sensitive to Parkinson’s disease and its treatment? NeuroImage 2010, 52:3. doi:10.1016/j.neuroimage.2009.12.080

Sadowska M, Sarecka-Hujar B, & Kopyta I. Cerebral Palsy: Current Opinions on Definition, Epidemiology, Risk Factors, Classification and Treatment Options. Neuropsychiatric disease and treatment 2020, 16. doi:10.2147/NDT.S235165

Sanger TD, Liker M, Arguelles E, et al. Pediatric Deep Brain Stimulation Using Awake Recording and Stimulation for Target Selection in an Inpatient Neuromodulation Monitoring Unit. Brain Sci 2018, 8:7. doi:10.3390/brainsci8070135

Schulz R, Wessel MJ, Zimerman M, et al. White Matter Integrity of Specific Dentato-Thalamo-Cortical Pathways is Associated with Learning Gains in Precise Movement Timing. Cerebral Cortex 2014, 25:7. doi:10.1093/cercor/bht356

Shakkottai VG, Batla A, Bhatia K, et al. Current Opinions and Areas of Consensus on the Role of the Cerebellum in Dystonia. Cerebellum (London, England) 2017, 16:2. doi:10.1007/s12311-016-0825-6

Solé-Padullés C, Castro-Fornieles J, de la Serna E, et al. Intrinsic connectivity networks from childhood to late adolescence: Effects of age and sex. Developmental Cognitive Neuroscience 2016, 17. doi:10.1016/j.dcn.2015.11.004

Winter S, Autry A, Boyle C, et al. Trends in the prevalence of cerebral palsy in a population-based study. Pediatrics 2002, 110:6. doi:10.1542/peds.110.6.1220

Zeidman P, Jafarian A, Seghier ML, et al. A guide to group effective connectivity analysis, part 2: Second level analysis with PEB. NeuroImage 2019, 200. doi:10.1016/j.neuroimage.2019.06.032

Zhang J, Li Z, Cao X, et al. Altered Prefrontal-Basal Ganglia Effective Connectivity in Patients With Poststroke Cognitive Impairment. Frontiers in neurology 2020, 11. doi:10.3389/fneur.2020.577482

