## Supplemental File A for "Dynamic Resting State Motor Network Connectivity of Neurotypical Children, the Groundwork for Network-Guided Therapy in Childhood Movement Disorders"

Supplementary Table A1. Left Hemisphere Connectivity Parameter Estimates and Confidence Intervals

|  |  | **Source** | | | | | | | | | | | | | | | | |
| --- | --- | --- | --- | --- | --- | --- | --- | --- | --- | --- | --- | --- | --- | --- | --- | --- | --- | --- |
|  |  | **L M1** | |  | **L STR** | |  | **L STN** | |  | **L GPi** | |  | **L THAL** | |  | **R CER** | |
|  |  | M | (95% CI) |  | M | (95% CI) |  | M | (95% CI) |  | M | (95% CI) |  | M | (95% CI) |  | M | (95% CI) |
| **Sink** | **L M1** | 0.39 | (0.34 , 0.43) |  | 0.30 | (0.26 , 0.34) |  | 0.09 | (0.07 , 0.12) |  | 0.07 | (0.04 , 0.11) |  | 0.10 | (0.06 , 0.14) |  | 0.00 | (0.00 , 0.00) |
|  | **L STR** | 0.00 | (0.00 , 0.00) |  | 0.47 | (0.42 , 0.52) |  | 0.11 | (0.09 , 0.14) |  | 0.00 | (0.00 , 0.00) |  | 0.12 | (0.07 , 0.16) |  | -0.10 | (-0.15 , -0.06) |
|  | **L STN** | -0.38 | (-0.44 , -0.33) |  | -0.35 | (-0.39 , -0.30) |  | -0.10 | (-0.14 , -0.06) |  | -0.17 | (-0.22 , -0.11) |  | -0.09 | (-0.14 , -0.05) |  | -0.47 | (-0.51 , -0.43) |
|  | **L GPi** | -0.19 | (-0.25 , -0.12) |  | 0.05 | (-0.01 , 0.11) |  | 0.07 | (0.04 , 0.09) |  | 0.16 | (0.12 , 0.20) |  | 0.13 | (0.10 , 0.17) |  | -0.18 | (-0.23 , -0.12) |
|  | **L THAL** | -0.17 | (-0.21 , 0.12) |  | 0.00 | (0.00 , 0.00) |  | 0.07 | (0.05 , 0.10) |  | -0.15 | (-0.20 , -0.11) |  | 0.22 | (0.17 , 0.26) |  | -0.20 | (-0.26 , -0.14) |
|  | **R CER** | 0.29 | (0.24 , 0.34) |  | 0.16 | (0.12 , 0.20) |  | 0.14 | (0.12 , 0.17) |  | 0.12 | (0.08 , 0.16) |  | 0.07 | (0.03 , 0.10) |  | 0.07* | (0.01 , 0.13) |

Parameter estimates for the Left Hemisphere model. All estimates are listed in Hz with the exception of self-connections. The self-connections are represented in the log scaling factor they are converted to during DCM inversion. This transformation ensures that all self-connections are negative when expressed in Hz, thus lending stability to the model. As such, more positive parameter estimates on self-connections indicate more self-inhibition and more negative (or less positive) parameter estimated on self-connections indicate less self-inhibition. The output from SPM reports self-connection parameter estimates in the log-scale, value, not Hz. (Note that, in the matrices in Fig. 1A and 1B, the diagonal has been converted to Hz using the formula A_Hz_ = -0.5 x exp(A) where A is the log-scaled parameter estimate value).

All non-zero parameter estimates had a posterior probability of 0.95 or greater with the exception of those with an asterisk. The right cerebellum self-connection had a posterior probability of 0.88 and the STR🡪GPi connection had a posterior probability of 0.75.

Supplementary Table A2. Right Hemisphere Connectivity Parameter Estimates and Confidence Intervals

|  |  | **Source** | | | | | | | | | | | | | | | | |
| --- | --- | --- | --- | --- | --- | --- | --- | --- | --- | --- | --- | --- | --- | --- | --- | --- | --- | --- |
|  |  |  | **R M1** |  |  | **R STR** |  |  | **R STN** |  |  | **R GPi** |  |  | **R THAL** |  |  | **L CER** |
|  |  | M | (95% CI) |  | M | (95% CI) |  | M | (95% CI) |  | M | (95% CI) |  | M | (95% CI) |  | M | (95% CI) |
| **Sink** | **R M1** | 0.25 | (0.21 , 0.30) |  | 0.02* | (-0.02 , 0.07) |  | 0.03* | (-0.01 , 0.07) |  | -0.06 | (-0.10 , -0.03) |  | 0.14 | (0.10 , 0.18) |  | 0.00 | (0.00 , 0.00) |
|  | **R STR** | 0.00 | (0.00 , 0.00) |  | 0.34 | (0.30 , 0.38) |  | 0.12 | (0.09 , 0.14) |  | 0.02* | (-0.02 , 0.05) |  | 0.00 | (0.00 , 0.00) |  | -0.18 | (-0.21 , -0.14) |
|  | **R STN** | -0.05* | (-0.12 , 0.03) |  | -0.26 | (-0.30 , -0.21) |  | -0.25 | (-0.29 , -0.21) |  | -0.24 | (-0.29 , -0.19) |  | -0.09 | (-0.14 , -0.04) |  | -0.45 | (-0.50 , -0.41) |
|  | **R GPi** | -0.03* | (-0.08 , 0.02) |  | -0.08 | (-0.12 , -0.04) |  | 0.04 | (0.02 , 0.07) |  | 0.08 | (0.03 , 0.12) |  | 0.00 | (0.00 , 0.00) |  | -0.08 | (-0.13 , -0.03) |
|  | **R THAL** | -0.09 | (-0.13 , -0.05) |  | 0.22 | (0.16 , 0.27) |  | 0.05 | (0.03 , 0.08) |  | 0.00 | (0.00 , 0.00) |  | 0.19 | (0.14 , 0.23) |  | -0.26 | (-0.31 , -0.21) |
|  | **L CER** | 0.09 | (0.04 , 0.13) |  | 0.24 | (0.20 , 0.29) |  | 0.08 | (0.05 , 0.11) |  | 0.00 | (0.00 , 0.00) |  | 0.05* | (-0.02 , 0.11) |  | -0.23 | (-0.30 , -0.16) |

Parameter estimates for the Right Hemisphere model. All estimates are listed in Hz with the exception of self-connections. The self-connections are represented in the log scaling factor they are converted to during DCM inversion. This transformation ensures that all self-connections are negative when expressed in Hz, thus lending stability to the model. As such, more positive parameter estimates on self-connections indicate more self-inhibition and more negative (or less positive) parameter estimated on self-connections indicate less self-inhibition. The output from SPM reports self-connection parameter estimates in the log-scale, value, not Hz. (Note that, in the matrices in Fig. 1A and 1B, the diagonal has been converted to Hz using the formula A_Hz_ = -0.5 x exp(A) where A is the log-scaled parameter estimate value).

All non-zero parameter estimates had a posterior probability of 0.95 or greater except for those with an asterisk. The posterior probabilities of those values are: M1🡪STN: 0.65; M1🡪GPi: 0.6; STR🡪M1: 0.56; STN🡪M1: 0.7; GPi🡪STR: 0.49; THAL🡪CER: 0.68.
