## Supplemental File B for "Dynamic Resting State Motor Network Connectivity of Neurotypical Children, the Groundwork for Network-Guided Therapy in Childhood Movement Disorders"

Supplementary Table B1. Left Hemisphere Connectivity association with (right hand) Purdue Pegboard Test score parameter estimates and confidence intervals

|  |  | **Source** | | | | | | | | | | | | | | | | |
| --- | --- | --- | --- | --- | --- | --- | --- | --- | --- | --- | --- | --- | --- | --- | --- | --- | --- | --- |
|  |  |  | **L M1** |  |  | **L STR** |  |  | **L STN** |  |  | **L GPi** |  |  | **L THAL** |  |  | **R CER** |
|  |  | M | (95% CI) |  | M | (95% CI) |  | M | (95% CI) |  | M | (95% CI) |  | M | (95% CI) |  | M | (95% CI) |
| **Sink** | **L M1** | 0.04 | (0.02 , 0.07) |  | 0.00 | (0.00 , 0.00) |  | -0.02 | (-0.04 , -0.01) |  | 0.00 | (0.00 , 0.00) |  | 0.04 | (0.02 , 0.06) |  | 0.00 | (0.00 , 0.00) |
|  | **L STR** | 0.05 | (0.03 , 0.07) |  | -0.06 | (-0.08 , -0.03) |  | 0.00 | (0.00 , 0.00) |  | 0.07 | (0.04 , 0.09) |  | 0.00 | (0.00 , 0.00) |  | 0.00 | (0.00 , 0.00) |
|  | **L STN** | 0.07 | (0.04 , 0.10) |  | -0.05 | (-0.08 , -0.03) |  | 0.01* | (-0.01 , 0.04) |  | 0.00 | (0.00 , 0.00) |  | 0.05 | (0.03 , 0.07) |  | 0.03 | (0.00 , 0.06) |
|  | **L GPi** | 0.06 | (0.03 , 0.09) |  | -0.07 | (-0.10 , -0.05) |  | 0.02* | (0.00 , 0.03) |  | 0.00 | (0.00 , 0.00) |  | -0.03 | (-0.05 , -0.01) |  | -0.04* | (-0.08 , -0.01) |
|  | **L THAL** | -0.07 | (-0.09 , -0.04) |  | -0.03* | (-0.07 , 0.02) |  | 0.00 | (0.00 , 0.00) |  | 0.05 | (0.03 , 0.07) |  | -0.04 | (-0.06 , -0.02) |  | 0.00* | (0.00 , 0.00) |
|  | **R CER** | 0.04 | (0.02 , 0.07) |  | 0.02* | (0.00 , 0.05) |  | -0.03 | (-0.04 , -0.01) |  | 0.00 | (0.00 , 0.00) |  | 0.04 | (0.03 , 0.06) |  | 0.00 | (0.00 , 0.00) |

Parameter estimates for the Left Hemisphere associated with right hand Purdue Pegboard Test (PPBT) score. For self-connections, a positive parameter estimate on a self-connection (positive effect of a covariate) indicates a positive relationship between the covariate and the level of self-inhibition and a negative parameter estimate on a self-connection (negative effect of a covariate) indicates a negative relationship between the covariate and the level of self-inhibition.

For all other connections, positive parameters indicate that more excitatory/less inhibitory connectivity values are associated with higher PPBT scores (positive relationship) and negative parameter values indicate that more inhibitory/less excitatory connectivity values are associated with higher PPBT scores (negative relationship).

All non-zero parameter estimates had a posterior probability of 0.95 or greater with the exception of those with an asterisk. The posterior probabilities of the asterisked values are: STR🡪THAL: 0.63; STR🡪CER: 0.75; STN🡪STN: 0.6; STN🡪GPi: 0.75; CER🡪GPi: 0.83; CER🡪THAL: 0.91.

Supplementary Table B2. Right Hemisphere Connectivity association with (left hand) Purdue Pegboard Test score parameter estimates and confidence intervals

|  |  | **Source** | | | | | | | | | | | | | | | | |
| --- | --- | --- | --- | --- | --- | --- | --- | --- | --- | --- | --- | --- | --- | --- | --- | --- | --- | --- |
|  |  |  | **R M1** |  |  | **R STR** |  |  | **RSTN** |  |  | **R GPi** |  |  | **R THAL** |  |  | **L CER** |
|  |  | M | (95% CI) |  | M | (95% CI) |  | M | (95% CI) |  | M | (95% CI) |  | M | (95% CI) |  | M | (95% CI) |
| **Sink** | **R M1** | 0.00 | (0.00 , 0.00) |  | -0.06 | (-0.08 , -0.04) |  | 0.00 | (0.00 , 0.00) |  | 0.00 | (0.00 , 0.00) |  | 0.00 | (0.00 , 0.00) |  | 0.00 | (0.00 , 0.00) |
|  | **R STR** | 0.00 | (0.00 , 0.00) |  | 0.10 | (0.08 , 0.13) |  | -0.06 | (-0.07 , -0.04) |  | -0.03 | (-0.05 , -0.01) |  | 0.07 | (0.05 , 0.08) |  | -0.04 | (-0.05 , -0.02) |
|  | **R STN** | 0.00 | (0.00 , 0.00) |  | 0.05 | (0.03 , 0.08) |  | 0.04 | (0.02 , 0.06) |  | 0.02* | (-0.02 , 0.06) |  | 0.06 | (0.03 , 0.08) |  | 0.01* | (-0.02 , 0.05) |
|  | **R GPi** | -0.04 | (-0.06 , -0.02) |  | 0.05 | (0.03 , 0.07) |  | -0.04 | (-0.06 , -0.02) |  | 0.00 | (0.00 , 0.00) |  | -0.05 | (-0.07 , -0.02) |  | -0.06 | (-0.08 , -0.03) |
|  | **R THAL** | -0.04 | (-0.06 , -0.02) |  | -0.05 | (-0.08 , -0.02) |  | 0.00 | (0.00 , 0.00) |  | 0.00 | (0.00 , 0.00) |  | -0.05 | (-0.07 , -0.03) |  | -0.07 | (-0.10 , -0.04) |
|  | **L CER** | -0.04 | (-0.06 , -0.01) |  | 0.00 | (0.00 , 0.00) |  | 0.03 | (0.02 , 0.05) |  | 0.00 | (0.00 , 0.00) |  | 0.11 | (0.08 , 0.14) |  | -0.13 | (-0.17 , -0.09) |

Parameter estimates for the Right Hemisphere associated with left hand Purdue Pegboard Test (PPBT) score. For self-connections, a positive parameter estimate on a self-connection (positive effect of a covariate) indicates a positive relationship between the covariate and the level of self-inhibition and a negative parameter estimate on a self-connection (negative effect of a covariate) indicates a negative relationship between the covariate and the level of self-inhibition.

For all other connections, positive parameters indicate that more excitatory/less inhibitory connectivity values are associated with higher PPBT scores (positive relationship) and negative parameter values indicate that more inhibitory/less excitatory connectivity values are associated with higher PPBT scores (negative relationship).

All non-zero parameter estimates had a posterior probability of 0.95 or greater with the exception of those with an asterisk. The posterior probabilities of the asterisked values are: GPi🡪STN: 0.57; CER🡪STN: 0.51.
