## Supplemental File C for "Dynamic Resting State Motor Network Connectivity of Neurotypical Children, the Groundwork for Network-Guided Therapy in Childhood Movement Disorders"

Supplementary Table C. Leave-one-out cross-validation results

| **Left Hemisphere** | | | |  | | **Right Hemisphere** | | |
| --- | --- | --- | --- | --- | --- | --- | --- | --- |
| Connection | *r*(17) | *p* |  | | Connection | | *r*(16) | *p* |
| M1→M1 | 0.15 | 0.26 |  | | M1→GPi | | 0.24 | 0.17 |
| **M1→Striatum** | **0.49** | **0.02** |  | | M1→Thalamus | | 0.11 | 0.33 |
| M1→STN | -0.27 | 0.86 |  | | M1→Cerebellum | | -0.19 | 0.78 |
| M1→GPi | -0.5 | 0.99 |  | | **Striatum→M1** | | **0.43** | **0.04** |
| **M1→Thalamus** | **0.43** | **0.03** |  | | Striatum→Striatum | | 0.28 | 0.13 |
| M1→Cerebellum | 0.15 | 0.28 |  | | Striatum→STN | | -0.1 | 0.65 |
| Striatum→Striatum | -0.06 | 0.6 |  | | Striatum→GPi | | -0.28 | 0.87 |
| Striatum→STN | -0.02 | 0.54 |  | | Striatum→Thalamus | | -0.13 | 0.7 |
| Striatum→GPi | 0.26 | 0.14 |  | | STN→Striatum | | 0.39 | 0.06 |
| STN→M1 | 0.03 | 0.45 |  | | STN→STN | | -0.15 | 0.72 |
| STN→Cerebellum | 0.19 | 0.21 |  | | **STN→GPi** | | **0.48** | **0.02** |
| GPi→Striatum | 0.3 | 0.1 |  | | STN→Cerebellum | | 0.06 | 0.4 |
| GPi→Thalamus | 0.28 | 0.12 |  | | GPi→Striatum | | -0.33 | 0.91 |
| Thalamus→M1 | 0.25 | 0.15 |  | | **Thalamus→Striatum** | | **0.46** | **0.03** |
| Thalamus→STN | 0.28 | 0.13 |  | | Thalamus→STN | | 0.23 | 0.18 |
| Thalamus→GPi | 0.19 | 0.22 |  | | Thalamus→GPi | | -0.33 | 0.93 |
| Thalamus→Thalamus | -0.32 | 0.91 |  | | Thalamus→Thalamus | | -0.16 | 0.74 |
| **Thalamus→Cerebellum** | **0.42** | **0.04** |  | | **Thalamus→Cerebellum** | | **0.58** | **0.006** |
|  |  |  |  | | Cerebellum→Striatum | | 0.07 | 0.39 |
|  |  |  |  | | Cerebellum→GPi | | -0.23 | 0.81 |
|  |  |  |  | | Cerebellum→Thalamus | | 0.11 | 0.33 |
|  |  |  |  | | Cerebellum→Cerebellum | | 0.18 | 0.23 |

Leave-one-out (LOO) cross-validation for covariation (Purdue Pegboard Testing score (mean-centered) for contralateral hand vs. connectivity estimate) connectivity estimates that had greater than 0.95 posterior probability are listed for each hemisphere. For each connection there is listed *r,* the Pearson out-of samples correlation coefficient value between the actual subject effect and the estimated subject effect as well as the p-value. Degrees of freedom are in the parentheses (17 for the left hemisphere, 16 for the right, due to difference in number of subjects in each hemisphere model). Connections with a p-value less than 0.05 are bolded (uncorrected).
